# Super-enhancers are transcriptionally more active and cell-type-specific than stretch enhancers

**DOI:** 10.1101/310839

**Authors:** Aziz Khan, Anthony Mathelier, Xuegong Zhang

**Affiliations:** Centre for Molecular Medicine Norway (NCMM), Nordic EMBL Partnership, University of Oslo, Oslo, Norway; Key Lab of Bioinformatics/Bioinformatics Division, BNRIST (Beijing National Research Center for Information Science and Technology), Department of Automation, Tsinghua University, Beijing 100084, China; Department of Cancer Genetics, Institute for Cancer Research, Oslo University Hospital Radiumhospitalet, 0372 Oslo, Norway; School of Life Sciences, Tsinghua University, Beijing, 100084, China

**Keywords:** Gene regulation, genomics, epigenomics, super-enhancers, stretch enhancers

## Abstract

**Background:** Super-enhancers and stretch enhancers represent classes of transcriptional enhancers that have been shown to control the expression of cell identity genes and carry disease- and trait-associated variants. Specifically, super-enhancers are clusters of enhancers defined based on the binding occupancy of master transcription factors (TFs), chromatin regulators, or chromatin marks, while stretch enhancers are large chromatin-defined regulatory regions of at least 3,000 base pairs. Several studies have characterized these regulatory regions in numerous cell types and tissues to decipher their functional importance. However, the differences and similarities between these regulatory regions have not been fully assessed.

**Results:** We integrated genomic, epigenomic, and transcriptomic data from ten human cell types to perform a comparative analysis of super and stretch enhancers with respect to their chromatin profiles, cell-type-specificity, and ability to control gene expression. We found that stretch enhancers are more abundant, more distal to transcription start sites, cover twice as much the genome and are significantly less conserved than super-enhancers. In contrast, super-enhancers are significantly more enriched for active chromatin marks and cohesin complex and transcriptionally active than stretch enhancers. Importantly, a vast majority of superenhancers (85%) overlap with only a small subset of stretch enhancers (13%), which are enriched for cell-type-specific biological functions, and control cell identity genes.

**Conclusions:** These results suggest that super-enhancers are transcriptionally more active and cell-type-specific than stretch enhancers, and importantly, most of the stretch enhancers that are distinct from superenhancers do not show an association with cell identity genes, are less active, and more likely to be poised enhancers.

## Background

The human body contains several hundred distinct cell types and most of the regulatory code that drives cell type-specific gene expression resides in *cis*-regulatory elements termed enhancers [1]. Enhancers are noncoding regulatory regions distal to the genes they regulate where transcription factors (TFs) and the transcriptional apparatus bind and orchestrate the gene expression regulation [1,2]. While estimations predicted approximately one million potential enhancers in the human genome, only a very small fraction of these enhancers are active in a given cell [3,4], marked by mono methylation of histone H3 at lysine 4 (H3K4me1) and acetylation of histone H3 at lysine 27 (H3K27ac) [5–8]. These active enhancers are primarily found in regions of accessible chromatin [5] and are significantly loaded with the coactivator protein p300 [9]. Other histone modification marks such as H3K79me3 as well as the occupancy of RNA Pol II delineate active enhancers [8]. Considerable levels of H3K4me3 are also observed at active enhancers bound by RNA Pol II [10]. Some of these enhancers are bi-directionally transcribed to produce RNA transcripts, referred to as enhancer RNAs (eRNAs) [11]. Additionally, studies have linked H3K4me3 broad peaks with transcriptional elongation, enhancer activity, and cellular identity [12,13]. Further, poised enhancers are enriched for trimethylation of histone H3 on lysine 27 (H3K27me3) and have depleted H3K27ac signal [6,14].

Despite critical advances in terms of technology and methodology to identify enhancers genome-wide and to understand their molecular mechanisms and function, a clear understanding is still lacking. Defining cell-type-specific enhancers and accurately assign them to the gene(s) they regulate is of great interest but very challenging due to lack of known cell type-specific signatures. In 2013, Whyte et al. showed that clusters of active enhancers, termed super-enhancers, control cell-identity. These super-enhancers were defined based on their enrichment for binding of key master regulator TFs, Mediator, and chromatin regulators [15]. These cluster of enhancers are cell type-specific, control the expression of cell-identity genes, are sensitive to perturbation, associated with disease, and boost the processing of primary microRNA into precursors of microRNAs [16,15,17,18]. Concomitantly to the discovery of super-enhancers, Parker et al. showed that large genomic regions with enhancer characteristic termed stretch enhancers, and defined based on their size (>3kb) control cell-identity [19]. These stretch enhancers are known to be cell-type-specific and are enriched for disease-associated variants, for instance carrying SNPs associated with type 2 diabetes [19,20]. Further, it has been shown that super- and stretch enhancers overlap with the known locus control regions (LCR) [21,16,19]. A significant attention has been given to these cell type-specific regulatory regions to understand their molecular mechanisms and functional importance. Since the parallel publications of these two concepts, there has been a confusion among some of the research community to differentiate these two classes of regulatory regions. It is unclear whether super- and stretch enhancers have different or equivalent regulatory potential. Some recent studies even referred to these regions collectively as SEs (Super/Stretch Enhancers), assuming them to be equivalent [22–24]. While some studies showed that super- and stretch enhancers overlap [25,26], however, a detailed comparative analysis to understand their differences and similarities is lacking.

We performed a comprehensive analysis of super- and stretch enhancers in 10 human cell-lines by integrating ChIP-seq data for histone modifications (H3K27ac, H3K4me1, H3K4me3, and H3K27me3), RNA Pol II, P300, cohesin components (RAD21 and SMC3) and CTCF, chromatin accessibility data (DNase-seq), and transcriptomics data (RNA-seq, GRO-seq, and GRO-cap). Our analyses revealed significant differences between super and stretch enhancers. Stretch enhancers are more abundant, distal to transcription start sites (TSS) and less conserved than super-enhancers. Comparatively, super-enhancers are significantly more enriched for active chromatin marks, RNA Pol II, and cohesin components, are transcriptionally more active, and transcribed than stretch enhancers. Finally, only a small fraction of stretch enhancers overlap with superenhancers, which we named super-stretch enhancers. These super-stretch enhancers are highly transcribed, associated with cell identity genes, and enriched for cell-type-specific biological functions.

## Results

### Genomic distribution and conservation of super- and stretch enhancers from ten human cell types

To systematically compare super- and stretch enhancers, we obtained super-enhancers from dbSUPER [27] and stretch enhancers from [19] from ten human cell types. These cell types included B-lymphoblastoid cells (GM128278), embryonic stem cells (H1-ES), erythrocytic leukaemia cells (K562), hepatocellular carcinoma cells (HepG2), human umbilical vein endothelial cells (HUVEC), human mammary epithelial cells (HMEC), human smooth muscle myoblasts (HSMM), normal human epidermal keratinocytes (NHEK), normal human lung fibroblasts (NHLF) and pancreatic islets (Islets). We looked at the genomic distribution of these regulatory regions and found that stretch enhancers are an order of magnitude more numerous and cover twice as much the human genome than super-enhancers (**Figure 1a, b**). Specifically, we found an average of 745 superenhancers with mean size 22,812 bp and 11,160 stretch enhancers with mean size 5,060 bp. Further, a majority of super-enhancers (69%) was located close to TSSs (<2kb), while a majority of stretch enhancers (70%) was located very distal from TSSs (>10kb) (**Figure 1c, Supplementary Figure S1**). Next, we investigated the evolutionary conservation of super- and stretch enhancers using the phastCons scores [28]. Superenhancers were significantly more conserved than stretch enhancers (p-value < 2.2e-16, Wilcoxon rank sum test) (**Figure 1d, S2**). Taken together, these results indicate that stretch enhancers are more distal to TSSs, more abundant in number and cover twice as much the genome than super-enhancers, which are significantly more conserved.

**Figure 1:**
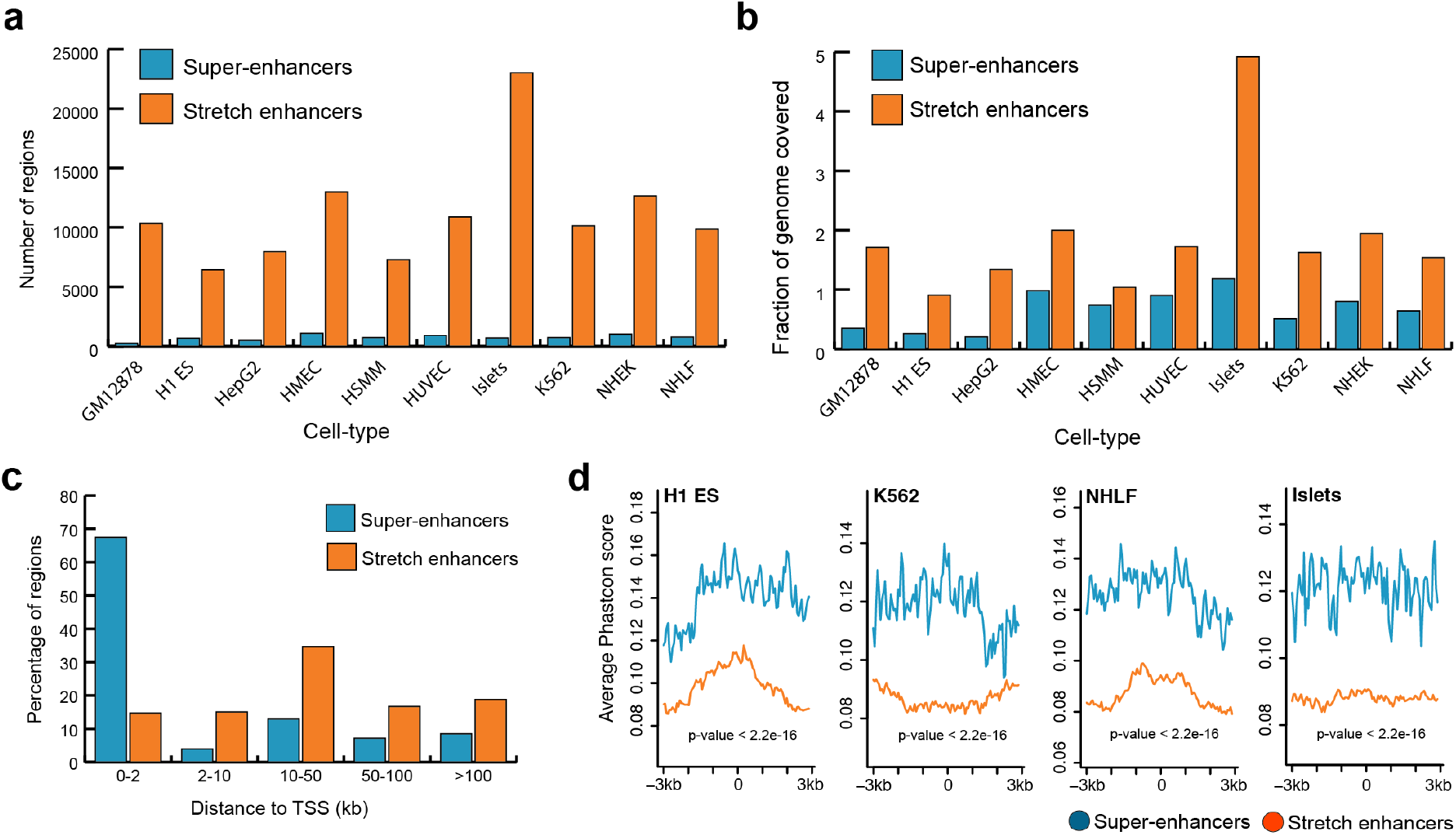
Genomic distribution and conservation of super- and stretch enhancers in 10 human cell types. (**a**) Number of super- and stretch enhancers in 10 cell types. (**b**) Fraction of the human genome covered by super- and stretch enhancers across 10 human cell types. (**c**) Distribution of distances to TSS for super- and stretch enhancers (average across the 10 cell types). (**d**) Evolutionary conservation score (phastCons) at super- and stretch enhancers with 6kb flanking regions in H1 -ES, K562, NHLF, and Islets cell types.

### Super-enhancers are enriched for active chromatin marks

We next sought to highlight the potentially distinct chromatin marks found at super- and stretch enhancers by using ChIP-seq data from the ENCODE project [29] for H3K27ac, H3K4me1, H3K4me3, and H3K27me3. Looking at average ChIP-seq signals, we observed that super-enhancers are highly enriched for active chromatin marks such as H3K27ac and H3K4me3, while depleted for poised marks such as H3K27me3 (**Figure 2a, b**). In contrast, stretch enhancers are highly enriched for poised chromatin mark and depleted for active chromatin marks (**Figure 2a, b**). Additionally, super- and stretch enhancers are almost equally marked by H3K4me1 (**Figure 2a, b**).

**Figure 2.**
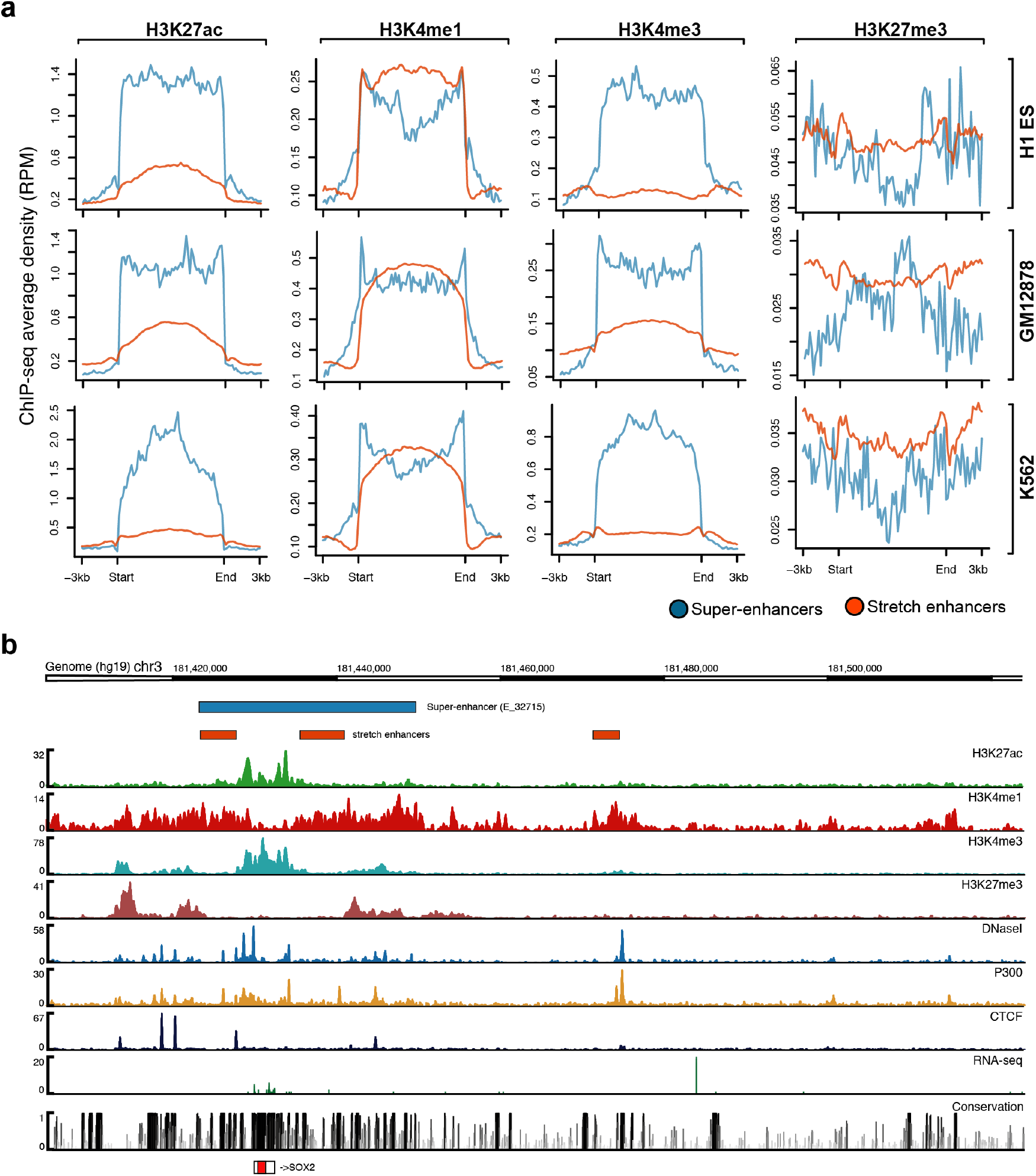
Chromatin modifications at super- and stretch enhancers. (**a**) Genome-wide average ChIP-seq profiles for H3K27ac, H3K4me, H3K4me3, and H3K27me3 at super- and stretch enhancers in H1-ESC, GM12878, and K562. (**b**) Genomic browser screenshot showing super- and stretch enhancers with ChIP-seq signals for H3K27ac, H3K4me1, H3K4me3, P300, and CTCF, and open chromatin (DNaseI), RNA-seq, and conservation at the locus of SOX2 gene in H1-ES cells.

Active enhancers are primarily found in regions of accessible chromatin [5]. Here, for most of the cell types, we observed significantly higher levels of DNase I hypersensitive sites (DHSs) for super-enhancers than for stretch enhancers (**Supplementary Figure S5**). Furthermore, we found that stretch enhancers overlap with super-enhancers at key cell-identity genes such as SOX2 (**Figure 2b**), POU5F/OCT4 (**Supplementary Figure S6A**), and NANOG (**Supplementary Figure S6B**) in embryonic stem cells, and are highly enriched for active chromatin marks. Additionally, in Pancreatic islet, super-enhancers have higher ChIP-seq signal for islet-specific transcription factors including PDX1, NKX2-2, FOXA2, and NKX6-1 [30] (**Supplementary Figure S7**).

Many type-2 diabetes SNPs from DIAGRAM (DIAbetes Genetics Replication And Meta-analysis) (red color) [31] as well as fasting glycemia SNPs from MAGIC (Meta-Analyses of Glucose and Insulin-related traits Consortium) (blue color) [32] were observed on and around these enhancer regions. Interestingly, we observed higher ChIP-seq binding signal at stretch enhancers that overlap with super-enhancers Taken together, these results highlight that super-enhancers are significantly more active and located in open regions than stretch enhancers, which are more likely to be poised.

### Super-enhancers are enriched with cohesin and CTCF binding

It is known that enhancers are brought close to their target genes through chromatin looping mechanisms. These long-range enhancer-promoter interactions and DNA looping are mediated by the cohesin complex and CTCF [33,34]. We compared the occupancy of two cohesin components (SMC3 and RAD21) and CTCF at super- and stretch enhancers in GM12878 and K562 cells. We observed significantly higher SMC3, RAD21, and CTCF ChIP-seq binding signal at super-enhancers than at stretch enhancers (**Figure 3**). This suggests that super-enhancers are regions with frequent interactions mediated by the cohesin complex and CTCF when compared to stretch enhancers.

**Figure 3:**
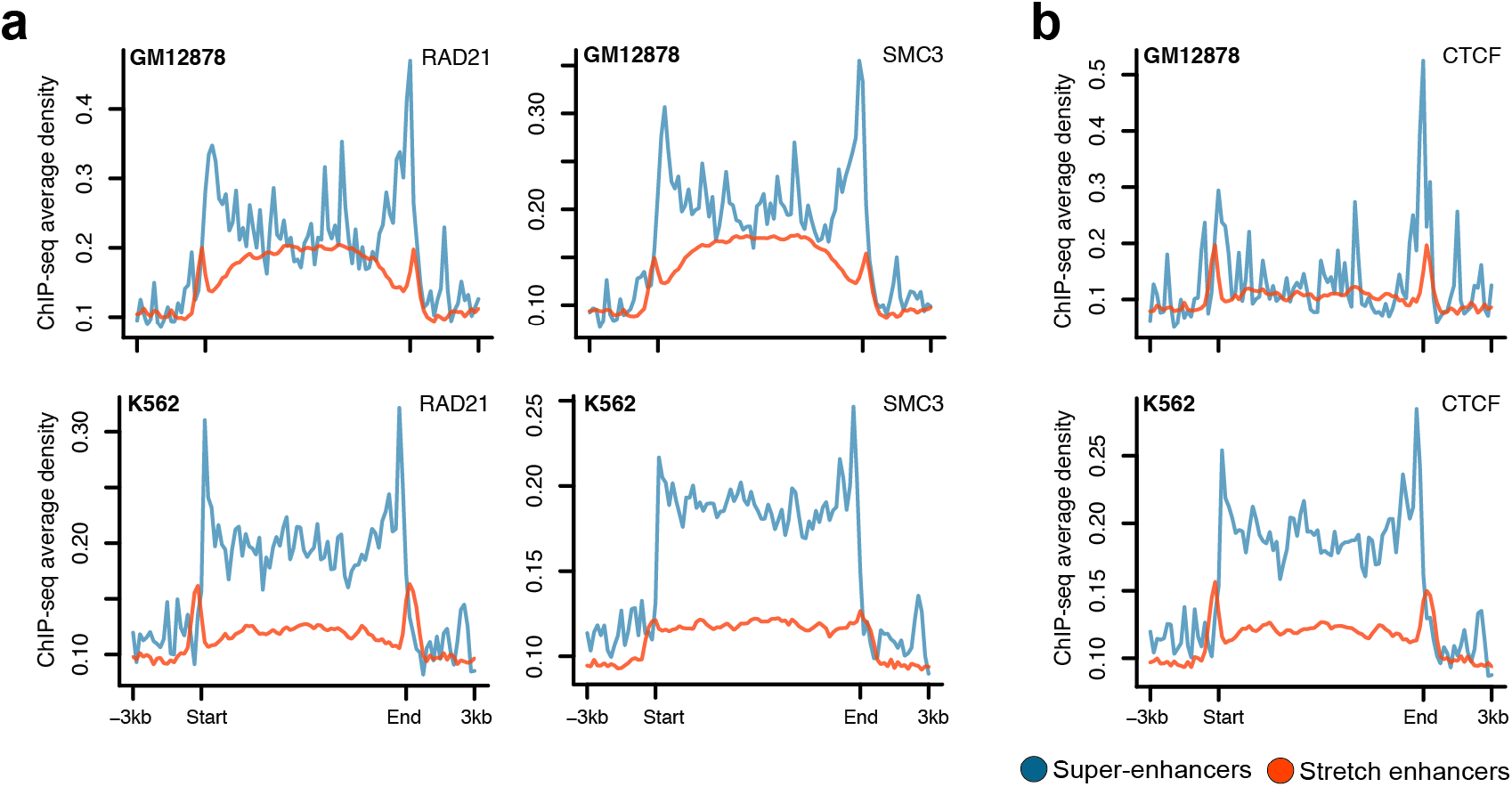
Chromatin organization at super- and stretch enhancers. (**a-b**) Spatial distribution of two Cohesin components RAD21 and SMC3 (**a**) and CTCF (**b**) at super- and stretch enhancers from K562 and GM12878 cells.

### Super-enhancers are transcriptionally more active than stretch enhancers

RNA Pol II plays a critical role in transcription and a majority of active enhancers recruit RNA Pol II [11]. We observed significantly higher RNA Pol II binding at super-enhancers than at stretch enhancers (**Figures 4a**). This was expected since we previously highlighted a significantly higher occupancy of active chromatin marks H3K27ac and H3K4me3 at super-enhancers. To further assess the effect of enhancer activity on gene expression regulation, we associated genes with super- and stretch enhancers based on proximity and calculated their transcriptional abundance. For all the 10 cell types, we found that the genes near superenhancers were significantly more expressed than genes near stretch-enhancers (p-value < 2.2e-16, Wilcoxon rank sum test) (**Figure 4b**).

**Figure 4.**
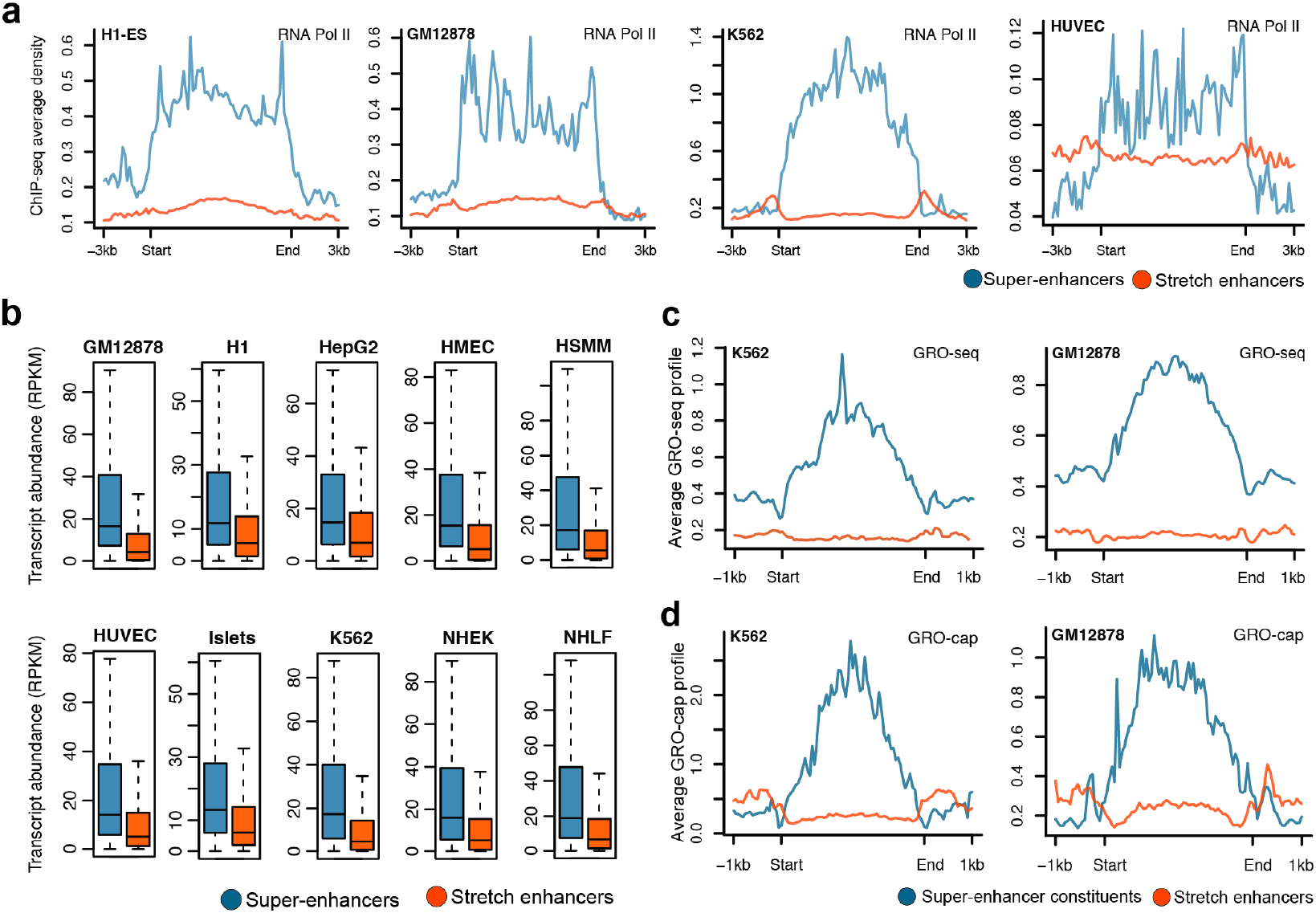
Transcriptional activity at super- and stretch enhancers. (**a**) Genome-wide profile of RNA Pol II at super- and stretch enhancers in H1-ESC, GM12878, K562, and HUVEC cell-lines. (**b**) Transcriptional abundance in reads per kilobase of transcript per million mapped reads (RPKM) of genes near super- and stretch enhancers across 10 cell types (p-value < 2.2e-16, Wilcoxon rank sum test). (**c**) GRO-seq profiles at the constituents of super- and stretch enhancers in K562 and GM12878 cell-lines. (**d**) GRO-cap profiles at the constituents of super- and stretch enhancers in K562 and GM1 2878 cell-lines.

Recent studies have shown that most of the active enhancers are bi-directionally transcribed to produce RNA transcripts, referred to as eRNAs [11]. These transcribed enhancers exhibit higher *in vitro* activity, suggesting that production of eRNA is linked to functional activity [35]. Recent techniques based on global run-on sequencing (GRO-seq and GRO-cap) have been developed for the detection of these unstable RNAs generated from enhancer elements [36,37]. We used publicly available data from GRO-seq and GRO-cap assays in K562 and GM12878 to investigate the levels of eRNAs at super- and stretch enhancers. Super-enhancers and their constituents (i.e. individual enhancers composing the clusters) harbored a significantly higher signal for both GRO-seq and GRO-cap (**Figure 4c-d, Supplementary Figure S8**) than stretch enhancers. Taken together, these results confirm that super-enhancers are more active and transcribed and can greatly enhance the transcription of their nearby genes when compared with stretch enhancers.

### A small subset of stretch enhancers overlap with super-enhancers

Next, we investigated to what extend super- and stretch enhancers are conserved between cell types and overlapped together. We computed pairwise Jaccard statistics between the cell types for super- and stretch enhancer regions, independently. We observed higher Jaccard statistics for stretch enhancers than for superenhancers, meaning that stretch enhancers were less cell-type-specific (Supplementary Figure S9). Next, we computed the fraction of overlap between super- and stretch enhancers in each cell type (**Figure 5a**). We observed that most of the super-enhancers overlap with a small fraction of the stretch enhancers in all cell types. On average, a vast majority of super-enhancers (85%) overlap with only a small number of stretch enhancers (13%) (**Figure 5b**). We termed the overlapping regions super-stretch enhancers. These super-stretch enhancers are significantly smaller in size than the remaining stretch enhancers (p-value < 2.2e-16, Wilcoxon rank sum test) (**Figure 5c**).

**Figure 5:**
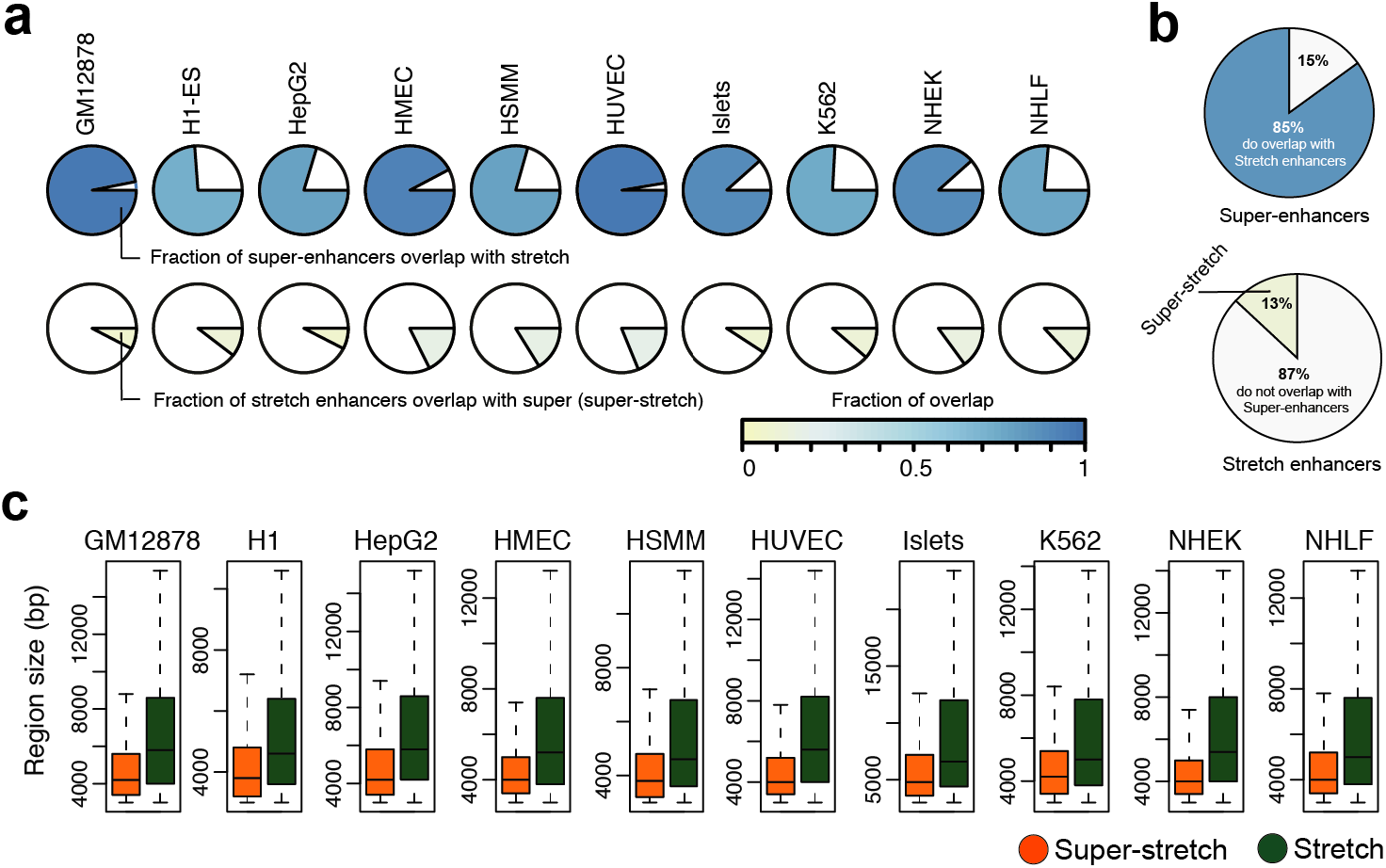
Overlap analysis of super- and stretch enhancers. (**a**) Fraction of overlap between super- and stretch enhancers across the ten cell types. (**b**) Pie chart of average overlap of super- and stretch enhancers. (**c**) The region length in base pairs (bp) of super-stretch and stretch enhancers across 10 cell types (p-value < 2.2e-16, Wilcoxon rank sum test).

In line with the fact that the vast majority of super-enhancers are super-stretch enhancers, we recapitulated the facts that these regions were (i) enriched for active chromatin marks such as H3K27ac and H3K4me3 (Supplementary Figure S10), (ii) enriched for cohesin and CTCF binding (Supplementary Figures S11 and S12a), (iii) near highly expressed genes (p-value<2.2e-16, Wilcoxon rank sum test, Supplementary Figure S12a), and (iv) significantly transcribed when compared to stretch enhancers (Supplementary Figure S12b).

Taken together, these results show that a vast majority of super-enhancers also contain a small fraction of stretch enhancers. Further, the small fraction of stretch enhancers that overlap with super-enhancers (called super-stretch enhancers) are highly enriched for active chromatin marks, highly transcribed and can greatly enhance the transcription of their associated genes.

### Super-stretch enhancers are cell-type-specific and control key cell identity genes

We sought to analyze how super-stretch enhancers were associated with cell-type-specific genes. We performed *k*-means clustering based on the H3K27ac histone modification at super, super-stretch, and stretch enhancers in five cell types (GM12878, K562, H1-ES, HepG2, and HUVEC). We observed cell-type-specific clusters for all the three groups, but significantly stronger cell-type-specific signal at super and super-stretch enhancers than at stretch enhancers (Figure 6a). It was expected to observe higher level of H3K27ac signal at super-enhancers as these were defined solely based on H3K27ac signal while stretch enhancers were defined using several other chromatin marks including H3K27ac. To test if the signal was coming from the way superenhancers were identified, we redefined stretch enhancers using same candidate enhancers defined using H3K27ac peaks in H1-ES, GM12878 and K562 cell types. We then divided these stretch enhancers into super-stretch and stretch enhancers, as described above. Interestingly, we found similar patterns for H3K27ac (**Supplementary Figure S15A**), GRO-seq signal (**Supplementary Figure S15B**), and also found cell-type-specific GO terms for super and super-stretch enhancers but not stretch enhancers (**Supplementary Figure S16**). This reinforces our observations that super- and stretch enhancers are intrinsically different.

When considering the closest genes to super- or super-stretch enhancers defined in H1-ES cells, we found that they were highly expressed in H1-ES cells but not in other cell types (**Figure 6b**). This observation holds for the 10 considered cell types (**Supplementary Figure S13**). On the contrary, we did not observe such a cell-type-specific behavior for genes close to stretch enhancers (**Supplementary Figure S13**). To confirm this cell-type-specific behavior associated to super- and stretch enhancers, we further assess the biological function of these regulatory regions, we used the tool GREAT to perform gene ontology enrichment analysis from the genes close to super, super-stretch, and stretch enhancers. In all cell types analyzed, super- and super-stretch enhancers were found close to genes enriched for corresponding cell-type-specific functions. For example, in ES cells, terms like stem cell development, stem cell differentiation and stem cell activation were enriched for genes associated with super and super-stretch enhancer (**Figure 5c; Supplementary Figure S14**).

**Figure 6.**
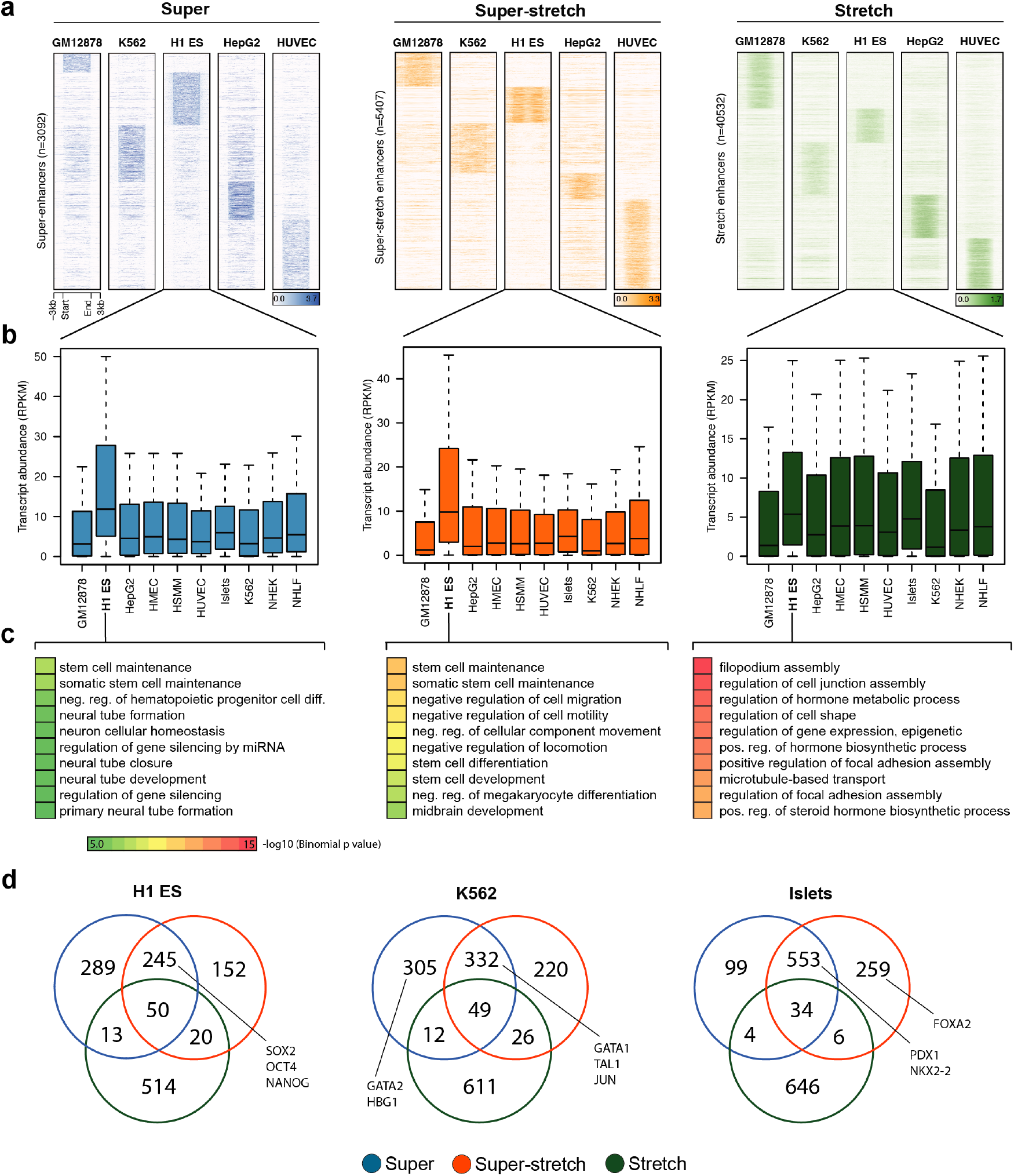
Cell-type-specificity analysis of super, super-stretch and stretch enhancers. (**a**) K-means clustering on the histone modification H3K27ac profile at super, super-stretch and stretch enhancers in five ENCODE cell types (GM12878, K562, H1-ES, HepG2, and HUVEC). (**b**) Transcriptional abundance in units of RPKM of genes associated with H1-ESC and how these genes are expressed in the other nine cell types tested as shown along the axis. (**c**) GO analysis of super, super-stretch and stretch enhancers in H1 ES cell type. (**d**) Venn diagram shows the overlap of genes associated with super, super-stretch and stretch enhancers, and label the known key cell-identity genes in H1 ES, K562, and Islets cells.

Next, we performed the overlap of genes associated with super-, super-stretch and stretch enhancers and checked for the known key cell-identity genes. We found a majority of key cell-identity genes associated with either super- or super-stretch enhancers. For example, the ESC pluripotency genes SOX2, OCT4 (POU5F1), and NANOG binding were found in super- and super-stretch enhancers but not stretch enhancers (**Figure 6d**). In K562 cells, proteins such as GATA1, TAL1, and JUN were binding at super- and super-stretch enhancers, while GATA2 and HBG1 at super-enhancers. Similarly, in Islets cells, PDX1 and NKX2-2 were binding at super- and super-stretch enhancers and FOXA2 at super-stretch enhancers. Taken together, these results suggest that a small subset of stretch enhancers, the one overlapping with super-enhancers (super-stretch), are preferentially associated with genes that have a key role in the cell-type-specific biology.

## Discussion

In this study, we have performed a comprehensive analysis of super- and stretch enhancers by comparing their histone modification profiles, chromatin accessibilities, and abilities to regulate cell-type-specific gene expression. At the genome scale, stretch enhancers are more abundant, cover twice the genome and are further away from TSS than super-enhancers; super-enhancers are evolutionary more conserved. Moreover, superenhancers are found to overlap active chromatin marks such as H3K27ac and H3K4me3 while stretch enhances are enriched for the poised mark H3K27me3. Super-enhancers are found to be significantly more occupied by the cohesin complex, CTCF, and RNA Pol II and produce eRNAs. Further, a majority of the superenhancers overlaps with a small fraction of stretch enhancers; the overlapping regions are more cell-type-specific and found near genes involved in cell maintenance, differentiation, and development.

We observed that super-enhancers are loaded with active chromatin marks such as H3K27ac and RNA Pol II and produce eRNAs, while stretch enhances are enriched for H3K27me3, depleted for H3K27ac and do not produce eRNA. These properties show that super-enhancers are transcriptionally active, while stretch enhancers are poised.

As super-enhancers were originally defined using H3K27ac signal, it was expected that they were containing higher level of H3K27ac signal when compared to stretch enhancers, which were defined based on multiple chromatin marks. Redefining stretch enhancers using solely the H3K27ac mark did not change our observations of the difference between super- and stretch. This reinforces the fact that super-enhancers are overall more active than stretch enhancers, which are likely poised enhancers. Further studies will be required to find the specific function of the genomic regions defined as stretch enhancers and that do not overlap superenhancers.

Conventionally, enhancers and promoters are regarded as distinct cis-regulatory elements and separated using enrichment for histone modifications H3K4me1/2 and H3K27ac for enhancers while H3K4me3 for promoters, but a growing number of evidence suggest that enhancers and promoters are very similar [38–40]. The exceptionally high enrichment for H3K27ac and RNA Pol II combined with H3K4me3 (a mark usually associated with promoters) at super-enhancer regions suggest that super-enhancers may also work as promoters to regulate their target gene(s) or these clusters may encompass promoter sequences. This is in line with the enrichment of GRO-seq and GRO-cap signal at these super-enhancer regions, which confirms that these regions initiate transcription.

Stretch enhancers were defined as large (>3Kb) linear genomic regions with specific chromatin marks [19] while super-enhancers were defined as clusters of enhancers enriched for active chromatin marks. We observed that a small fraction of stretch enhancers that overlap super-enhancers was more active, cell-type-specific, and significantly smaller than the rest of these regions. It clearly highlights that length is not an optimal feature to determine cell-type-specific regulatory regions. Our analyses and several other lines of evidence suggest that enhancers are more likely to be cell-type-specific, transcriptionally active, and frequently interacting when found in clusters at the genomic scale, whatever their sizes [30,41]. Whether the individual enhancers within a cluster (super-enhancer) work synergistically or additively is still in active debate [42]. Some studies have showed synergy and hierarchy between the individual enhancers within the clusters [23,43,44], but this synergy is less obvious for some developmentally regulated super-enhancers [45]. Further, locus control regions [21] have been known for decades and whether the newly introduced super- and stretch enhancers represent a new paradigm in transcriptional regulation is still debated [26]. It has been suggested that genes that are regulated by multiple cis-regulatory elements may achieve higher levels of RNA Pol II recruitment due to higher local concentration of required factors than genes regulated by fewer regulatory elements [38,39]. A very recent simulation study, based on the properties of super-enhancers, proposed a conceptual phase separation model for transcriptional gene regulation [46]. If that is the case, then the current approaches to identify clusters of enhancers is not optimal. Indeed, the current methods do not integrate chromatin interaction data to restrict these clusters to be within the so-called topologically associated domains, where most of the enhancer-promoter interactions and regulation taking place. Hence, we still need to develop new methods to capture cell type-specific enhancers that might not be derived from clusters of enhancers at the genomic scale but rather in the 3D space of the nucleus.

## Material and methods

### Data description

We used the processed ChIP-seq data for histone modifications and DNase-seq data generated by ENCODE (Encyclopedia of Regulatory DNA Elements) to perform all the analysis in this report (hgdownload-test.cse.ucsc.edu/goldenPath/hg19/encodeDCC/) (**Table S1**). We obtained the processed RNA-seq based gene expression data as RPKM for the ten cell types from [19]. We also used GRO-seq and GRO-cap data in GM12878 and K562, which is listed in supplementary **Table S2**.

### GRO-seq and GRO-cap data analysis

The GRO-seq, and GRO-cap reads were aligned to human genome-build hg19 using bowtie2 [47] with default parameters. The aligned and sorted bam files were used to compute the average signal profile at superenhancer and stretch enhancers.

### Super-enhancers

Super-enhancers were downloaded as BED files for the ten human cell types GM12878, H1-ES, K562, HepG2, HUVEC, HMEC, HSMM, NHEK, NHLF, and Islets from dbSUPER (http://asntech.org/dbsuper/) [27].

### Stretch enhancers

We obtained the stretch enhancer annotations for the 10 human cell types (including GM12878, H1-ES, K562, HepG2, HUVEC, HMEC, HSMM, NHEK, NHLF and Islets) from [19]. We also redefined stretch enhancers using only H3K27ac ChIP-seq data in cell-lines H1-ES, GM12878, and K562. We used the stitched enhancers regions (typical enhancers and super-enhancers) and ranked them based on length and separated the one which are larger than 3kb.

### Assigning genes to super and stretch enhancers

Genes were assigned to super- and stretch enhancers based on proximity as descried in [15,16]. It is known that enhancers tend to loop and communicate with target genes [48], and most of these enhancer-promoter interactions occur within a distance of ~50kb [49]. This approach identified a large proportion of true enhancer/promoter interactions in ESC [50]. Hence, all transcriptionally active genes were assigned to superenhancers and stretch enhancers within a 50kb window.

### Gene ontology analysis

To perform Gene Ontology analysis, we used GREAT (Genomic Regions Enrichment of Annotations Tool) (http://bejerano.stanford.edu/great/, version 3.0.0) [51] with default parameters. The top 10 GO terms with the lowest p-value were reported.

### Overlap analysis

We used BEDTools [52] to perform intersection of genomic regions. We considered two regions to overlap if they shared at least 1 bp. To perform pairwise overlap analysis we used the Intervene tool [53].

### Visualization and statistical analysis

We generated box plots using the R programming language by extending the whiskers to 1.5x the interquartile range. The P-values for box-plots were calculated using Wilcoxon signed-rank tests with the wilcox.test function in R. We used ngs.plot [54] to generate heat maps and normalized binding profiles at the constituents of super- and stretch enhancers along with their flanking 3kb regions. We used Intervene (https://asntech.shinyapps.io/intervene/) [53] to generate pariwise heatmaps. For genome-browser screenshots, we used the Biodalliance genome browser with ENCODE data [55].

## List of abbreviations

TFs: : Transcription Factors;
ChIP-seq: : Chromatin immune precipitation followed by high-throughput sequencing;
ES: : Embryonic Stem;
SEs: : Super-enhancers;
LCR: : Locus Control Regions;
TSS: : Transcription Start Site;
GO: : Gene Ontology;
GRO-seq: : Global run-on sequencing;
GRO-cap: : Global run-on sequencing coupled with enrichment for nascent RNAs with 5’caps;
ENCODE: : Encyclopedia of Regulatory DNA Elements

## Declarations

### Authors’ contributions

AK conceived the project and designed the experiments. XG reviewed and approved experiment design and supervised the project. AM provided suggestions, supported to redefine stretch enhancers, and overviewed the finalization of the project. AK performed the experiments and analyzed the data. AK wrote the manuscript and XG and AM reviewed it. All authors read and approved the final manuscript.

## Acknowledgements

We thank the ENCODE consortium and all the authors of the original studies used in this work for making their data freely available.

## References

1. Heinz S, Romanoski CE, Benner C, Glass CK. The selection and function of cell type-specific enhancers. Nat Rev Mol Cell Biol. 2015;16:144–154.

2. Barolo S. Shadow enhancers: Frequently asked questions about distributed cis-regulatory information and enhancer redundancy. BioEssays. 2012;34:135–141.

3. Thurman RE, Rynes E, Humbert R, Vierstra J, Maurano MT, Haugen E, et al. The accessible chromatin landscape of the human genome. Nature. Nature Publishing Group; 2012;489:75–82.

4. Heintzman ND, Hon GC, Hawkins RD, Kheradpour P, Stark A, Harp LF, et al. Histone modifications at human enhancers reflect global cell-type-specific gene expression. Nature. Nature Publishing Group; 2009;459:108–12.

5. Shlyueva D, Stampfel G, Stark A. Transcriptional enhancers: from properties to genome-wide predictions. Nat Rev Genet. Nature Publishing Group; 2014;15:272–86.

6. Rada-Iglesias A, Bajpai R, Swigut T, Brugmann SA, Flynn RA, Wysocka J. A unique chromatin signature uncovers early developmental enhancers in humans. Nature. Nature Publishing Group; 2011;470:279–83.

7. Arnold CD, Gerlach D, Stelzer C, Boryń ŁM, Rath M, Stark A. Genome-wide quantitative enhancer activity maps identified by STARR-seq. Science. 2013;339:1074–7.

8. Bonn S, Zinzen RP, Girardot C, Gustafson EH, Perez-Gonzalez A, Delhomme N, et al. Tissue-specific analysis of chromatin state identifies temporal signatures of enhancer activity during embryonic development. Nat Genet. 2012;44:148–56.

9. Visel A, Blow MJ, Li Z, Zhang T, Akiyama JA, Holt A, et al. ChIP-seq accurately predicts tissue-specific activity of enhancers. Nature. 2009;457:854–8.

10. Calo E, Wysocka J. Modification of enhancer chromatin: what, how, and why? Mol Cell. Elsevier Inc.; 2013;49:825–37.

11. Kim T-K, Hemberg M, Gray JM, Costa AM, Bear DM, Wu J, et al. Widespread transcription at neuronal activity-regulated enhancers. Nature. 2010;465:182–187.

12. Chen K, Chen Z, Wu D, Zhang L, Lin X, Su J, et al. Broad H3K4me3 is associated with increased transcription elongation and enhancer activity at tumor-suppressor genes. Nat Genet. 2015;47:1149–57.

13. Benayoun BA, Pollina EA, Ucar D, Mahmoudi S, Karra K, Wong ED, et al. H3K4me3 breadth is linked to cell identity and transcriptional consistency. Cell. 2014;158:673–88.

14. Creyghton MP, Cheng AW, Welstead GG, Kooistra T, Carey BW, Steine EJ, et al. Histone H3K27ac separates active from poised enhancers and predicts developmental state. Proc Natl Acad Sci U S A. 2010;107:21931–6.

15. Whyte WA, Orlando DA, Hnisz D, Abraham BJ, Lin CY, Kagey MH, et al. Master Transcription Factors and Mediator Establish Super-Enhancers at Key Cell Identity Genes. Cell. Elsevier Inc.; 2013;153:307–19.

16. Hnisz D, Abraham BJ, Lee TI, Lau A, Saint-André V, Sigova AA, et al. Super-enhancers in the control of cell identity and disease. Cell. 2013;155:934–47.

17. Lovén J, Hoke HA, Lin CY, Lau A, Orlando DA, Vakoc CR, et al. Selective inhibition of tumor oncogenes by disruption of super-enhancers. Cell. 2013;153:320–34.

18. Suzuki HI, Young RA, Sharp PA. Super-Enhancer-Mediated RNA Processing Revealed by Integrative MicroRNA Network Analysis. Cell. 2017;168:1000–1014.e15.

19. Parker SCJ, Stitzel ML, Taylor DL, Orozco JM, Erdos MR, Akiyama JA, et al. Chromatin stretch enhancer states drive cell-specific gene regulation and harbor human disease risk variants. Proc Natl Acad Sci U S A. 2013;110:17921–6.

20. Quang DX, Erdos MR, Parker SCJ, Collins FS. Motif signatures in stretch enhancers are enriched for disease-associated genetic variants. Epigenetics Chromatin. 2015;8:23.

21. Li Q, Peterson KR, Fang X, Stamatoyannopoulos G. Locus control regions. Blood. 2002;100:3077–3086.

22. Vahedi G, Kanno Y, Furumoto Y, Jiang K, Parker SCJ, Erdos MR, et al. Super-enhancers delineate disease-associated regulatory nodes in T cells. Nature. 2015;520:558–62.

23. Hnisz D, Schuijers J, Lin CY, Weintraub AS, Abraham BJ, Lee TI, et al. Convergence of Developmental and Oncogenic Signaling Pathways at Transcriptional Super-Enhancers. Mol Cell. Elsevier Inc.; 2015;58:362–70.

24. Flynn RA, Do BT, Rubin AJ, Calo E, Lee B, Kuchelmeister H, et al. 7SK-BAF axis controls pervasive transcription at enhancers. Nat Struct Mol Biol. Nature Publishing Group; 2016;1–11.

25. Niederriter A, Varshney A, Parker S, Martin D. Super Enhancers in Cancers, Complex Disease, and Developmental Disorders. Genes. 2015;6:1183–200.

26. Pott S, Lieb JD. What are super-enhancers? Nat Genet. Nature Publishing Group; 2014;47:8–12.

27. Khan A, Zhang X. dbSUPER: a database of super-enhancers in mouse and human genome. Nucleic Acids Res. 2016;44:D164–D171.

28. Rosenbloom KR, Armstrong J, Barber GP, Casper J, Clawson H, Diekhans M, et al. The UCSC Genome Browser database: 2015 update. Nucleic Acids Res. 2014;43:D670–D681.

29. Dunham I, Kundaje A, Aldred SF, Collins PJ, Davis C a., Doyle F, et al. An integrated encyclopedia of DNA elements in the human genome. Nature. 2012;489:57–74.

30. Pasquali L, Gaulton KJ, Rodríguez-Seguí SA, Mularoni L, Miguel-Escalada I, Akerman I, et al. Pancreatic islet enhancer clusters enriched in type 2 diabetes risk-associated variants. Nat Genet. 2014;46:136–43.

31. Morris AP, Voight BF, Teslovich TM, Ferreira T, Segrè AV, Steinthorsdottir V, et al. Large-scale association analysis provides insights into the genetic architecture and pathophysiology of type 2 diabetes. Nat Genet. Nature Publishing Group, a division of Macmillan Publishers Limited. All Rights Reserved.; 2012;44:981–90.

32. Scott RA, Lagou V, Welch RP, Wheeler E, Montasser ME, Luan J, et al. Large-scale association analyses identify new loci influencing glycemic traits and provide insight into the underlying biological pathways. Nat Genet. Nature Publishing Group, a division of Macmillan Publishers Limited. All Rights Reserved.; 2012;44:991–1005.

33. Rao SSP, Huang S-C, Hilaire BGS, Engreitz JM, Perez EM, Kieffer-Kwon K-R, et al. Cohesin Loss Eliminates All Loop Domains. Cell. 2017;171:305–320.e24.

34. Zuin J, Dixon JR, van der Reijden MIJA, Ye Z, Kolovos P, Brouwer RWW, et al. Cohesin and CTCF differentially affect chromatin architecture and gene expression in human cells. Proc Natl Acad Sci USA. 2014;111:996–1001.

35. Andersson R, Gebhard C, Miguel-Escalada I, Hoof I, Bornholdt J, Boyd M, et al. An atlas of active enhancers across human cell types and tissues. Nature. 2014;507:455–61.

36. Core LJ, Waterfall JJ, Lis JT. Nascent RNA Sequencing Reveals Widespread Pausing and Divergent Initiation at Human Promoters. Science. 2008;322:1845–8.

37. Kruesi WS, Core LJ, Waters CT, Lis JT, Meyer BJ. Condensin controls recruitment of RNA polymerase II to achieve nematode X-chromosome dosage compensation. eLife. 2013;2:e00808.

38. Andersson R. Promoter or enhancer, what’s the difference? Deconstruction of established distinctions and presentation of a unifying model. BioEssays. 2015;37:314–23.

39. Andersson R, Sandelin A, Danko CG. A unified architecture of transcriptional regulatory elements. Trends Genet. Elsevier Ltd; 2015;31:426–33.

40. Kim T, Shiekhattar R. Architectural and Functional Commonalities between Enhancers and Promoters. Cell. Elsevier Inc.; 2015;162:948–59.

41. Ernst J, Kheradpour P, Mikkelsen TS, Shoresh N, Ward LD, Epstein CB, et al. Mapping and analysis of chromatin state dynamics in nine human cell types. Nature. 2011;473:43–9.

42. Dukler N, Gulko B, Huang Y, Siepel A. Is a super-enhancer greater than the sum of its parts? 2017;49:2–7.

43. Proudhon C, Snetkova V, Raviram R, Lobry C, Badri S, Jiang T, et al. Active and Inactive Enhancers Cooperate to Exert Localized and Long-Range Control of Gene Regulation. Cell Rep. 2016;15:2159–69.

44. Shin HY, Willi M, Yoo KH, Zeng X, Wang C, Metser G, et al. Hierarchy within the mammary STAT5-driven Wap super-enhancer. Nat Genet. 2016;48:904–11.

45. Hay D, Hughes JR, Babbs C, Davies JOJ, Graham BJ, Hanssen LLP, et al. Genetic dissection of the α-globin superenhancer in vivo. Nat Genet. 2016;1–12.

46. Hnisz D, Shrinivas K, Young RA, Chakraborty AK, Sharp PA. A Phase Separation Model for Transcriptional Control. Cell. 2017;169:13–23.

47. Langmead B, Trapnell C, Pop M, Salzberg SL. Ultrafast and memory-efficient alignment of short DNA sequences to the human genome. Genome Biol. 2009;10:R25.

48. Ong C-T, Corces VG. Enhancer function: new insights into the regulation of tissue-specific gene expression. Nat Rev Genet. Nature Publishing Group; 2011;12:283–93.

49. Chepelev I, Wei G, Wangsa D, Tang Q, Zhao K. Characterization of genome-wide enhancer-promoter interactions reveals co-expression of interacting genes and modes of higher order chromatin organization. Cell Res. Nature Publishing Group; 2012;22:490–503.

50. Dixon JR, Selvaraj S, Yue F, Kim A, Li Y, Shen Y, et al. Topological domains in mammalian genomes identified by analysis of chromatin interactions. Nature. Nature Publishing Group; 2012;485:376–80.

51. McLean CY, Bristor D, Hiller M, Clarke SL, Schaar BT, Lowe CB, et al. GREAT improves functional interpretation of cis-regulatory regions. Nat Biotechnol. Nature Publishing Group; 2010;28:495–501.

52. Quinlan AR, Hall IM. BEDTools: a flexible suite of utilities for comparing genomic features. Bioinforma Oxf Engl. 2010;26:841–2.

53. Khan A, Mathelier A. Intervene: a tool for intersection and visualization of multiple gene or genomic region sets. BMC Bioinformatics. 2017;18:287.

54. Shen L, Shao N, Liu X, Nestler E. ngs.plot: Quick mining and visualization of next-generation sequencing data by integrating genomic databases. BMC Genomics. BMC Genomics; 2014;15:284.

55. Down TA, Piipari M, Hubbard TJP. Dalliance: interactive genome viewing on the web. Bioinformatics. 2011;27:889–90.

